# Cargo crowding at actin-rich regions along axons causes local traffic jams in neurons

**DOI:** 10.1101/151761

**Authors:** Parul Sood, Kausalya Murthy, T. Vinod Kumar, Michael L Nonet, Gautam I. Menon, Sandhya P. Koushika

**Affiliations:** DBS-TIFR, HomiBhabha Road, Mumbai-400005, India; Neurobiology, NCBS-TIFR, Bellary Road, Bangalore-560065, India; The Institute of Mathematical Sciences, CIT Campus, Taramani, Chennai-600113, India; Department of Anatomy and Neurobiology, Washington University School of Medicine, St. Louis, MO 63110, USA; Homi Bhabha National Institute, Training School complex, Anushaktinagar, Mumbai-400094, India

## Abstract

Steady axonal cargo flow is central to the functioning of healthy neurons. However, a substantial fraction of cargo in axons remains stationary across a broad distribution of times. We examine the transport of pre-synaptic vesicles (pre-SVs), endosomes and mitochondria in *C. elegans* touch receptor neurons (TRNs), showing that stalled cargo are predominantly present at actin-rich regions along the neuronal process. Cargo stalled at actin-rich regions increase the propensity of moving cargo to stall at the same location, resulting in traffic jams. Such local traffic jams at actin-rich regions are likely to be a general feature of axonal transport since they occur in *Drosophila* neurons as well. These traffic jams can act as both sources and sinks of vesicles. We propose that they act as functional reservoirs that contribute to maintaining robust cargo flow in the neuron.

## INTRODUCTION

Microtubule-based molecular motors transport diverse neuronal cargo along the axon, both in the anterograde direction, away from the cell body towards the synapse, as well as in the retrograde direction towards the cell body. Time-lapse fluorescence imaging of *in vivo* cargo transport shows that mobile and stationary cargo co-exist along the axon (Kang, Tian et al. 2008, Tang, Scott et al. 2012, Tang, Scott et al. 2013, Iacobucci, Rahman et al. 2014). Although a significant fraction of vesicular cargo and mitochondria in neurons remains stationary over imaging time, studies thus far have largely focused on understanding the characteristics, mechanisms of motion and functions of moving cargo (Vale 2003, Hirokawa and Takemura 2004, Hirokawa and Takemura 2005, Hirokawa and Noda 2008, Hirokawa, Niwa et al. 2010, Maday, Twelvetrees et al. 2014, Gibbs, Greensmith et al. 2015). Moving vesicular cargo typically pause for up to 15 seconds due to motor based mechanisms (Conway, Wood et al. 2012, Gu, Sun et al. 2012, Bálint, Verdeny Vilanova et al. 2013). Cargo are also known to pause for up to several minutes (Kang, Tian et al. 2008, Tang, Scott et al. 2012, Tang, Scott et al. 2013, Iacobucci, Rahman et al. 2014, Sheng 2014) but we know little about the factors that cause cargo to remain immobile over these much longer timescales.

Mechanisms such as a tug-of-war between motors and motor accumulations at the ends of microtubules have been implicated in stalling of cargo *in vitro* (Vershinin, Carter et al. 2007, Dixit, Ross et al. 2008, Ross, Shuman et al. 2008, Conway, Wood et al. 2012, Leduc, Padberg-Gehle et al. 2012). The microtubule binding protein, Syntaphilin, has also been implicated in docking mitochondria to microtubules for up to several minutes (Kang, Tian et al. 2008). Such studies suggest that microtubule-associated proteins and motor-based mechanisms both play roles in interrupting smooth cargo transport. However, *in vitro* studies do not address the complexity of cellular environments where multiple mechanisms may contribute to stalling, nor is it clear if mechanisms of stalling depend on cargo or neuron type.

Constraints common to all types of moving cargo in axons include the narrow geometry of the axon and its cytoskeletal architecture. The axon is densely packed with microtubules of variable lengths, actin, intermediate filaments, cytoskeleton associated proteins as well as stalled cargo (Hirokawa 1982, Black and Kurdyla 1983, Cueva, Mulholland et al. 2007, Kaasik, Safiulina et al. 2007, Jaworski, Hoogenraad et al. 2008, Leterrier, Vacher et al. 2011, Xu, Zhong et al. 2013, Arnold and Gallo 2014, Narayanareddy, Vartiainen et al. 2014, Richardson, Spilker et al. 2014, Yau, van Beuningen et al. 2014, Coles and Bradke 2015, Ganguly, Tang et al. 2015, Che, Chowdary et al. 2016). An *in vivo* study suggests that synaptic vesicles tend to stop at microtubule ends due to the absence of a track to move ahead (Yogev, Cooper et al. 2016). Other *in vitro* studies, performed with motors and microtubules alone, show that crowding agents such as PEG can affect motor movement, as can crowding caused by motor accumulation at the end of the track (Leduc, Padberg-Gehle et al. 2012, L Conway and J L Ross 2014). Additionally, cargo sizes are varied, ranging from 30nm-3μm (Hirokawa 1982, Nakata, Terada et al. 1998, Cueva, Mulholland et al. 2007), and their motion is likely further constrained in cytoskeletally crowded environments in comparison to motors unbound to cargo. Since a network of filamentous actin is present under the plasma membrane in addition to deep actin along the axon (Xu, Zhong et al. 2013, Ganguly, Tang et al. 2015), actin enriched regions could be a potential source of such impediments to cargo motion. Many stationary endosomes have also been shown to be present at actin-rich regions in cultured hippocampal neurons (Ganguly, Tang et al. 2015).

In this study, we show that actin-rich regions along the axon act as hotspots that initiate local traffic jams by stalling moving cargo. We suggest that vesicles trapped at these local traffic jams may form functional reservoirs along the neuronal process and that such dynamic reservoirs may underlie the robustness of cargo transport within the crowded environment of the axon.

## RESULTS

### Multiple vesicular cargo in neurons are immobile for long durations

We tracked fluorescently labeled precursors of synaptic vesicles (pre-SVs) and endosomes using time-lapse fluorescence imaging in neurons of *C. elegans.* We used the following markers: (i) RAB-3 and Synaptobrevin-1 (SNB-1) for pre-SVs in TRNs, (ii) RAB-5 for endosomes in TRNs, and (iii) RAB-3 for pre-SVs in commissures of motor neurons.

Each cargo in a defined neuron has distinct movement parameters (Table S1). We label a tagged vesicle as stationary if it remains immobile for at least five times the maximum pause duration of its cognate moving vesicle in the neuron-type imaged (Table S2). This cut-off was chosen since we wished to exclude vesicles stalled due to either motor pausing or stalled as a consequence of a tug-of-war between oppositely directed motors. Both these processes typically occur at times scales of 0.5s-15s (Conway, Wood et al. 2012, Gu, Sun et al. 2012). The timescales of pausing could additionally depend on the nature of the cargo motor complex, the type of cargo, the size of the cargo and the axoplasmic environment of the neuron where these cargo move. We therefore independently calculated cut-offs for different cargo types and neuron types based on our measured pause times for each cargo type imaged (Table S1 and S2). We consistently observe stationary vesicles in all cargo types mentioned above (Figure 1A-E). These are seen in kymographs as long vertical lines (Figure 1C-E, red and green arrows). In TRNs of *C. elegans*, stationary pre-SVs are observed irrespective of the type of pre-SV protein tagged (RAB-3 or SNB-1), their position in the axon or the dosage of the marker (Figure S1A-G). We further split stationary cargo (SC) into two categories: short lived stationary cargo (sl) that are stationary for five times the maximum pause time (green arrows, Figure 1C-E, Table S2) and long-lived SC (ll) (red arrows, Figure 1C-E) that are stationary for three times the maximum stall time of a short-lived SC (Table S2). The proportions of short-lived and long-lived stationary pre-SVs marked by GFP::RAB-3 are 52% and 48% respectively. There are typically about five locations where long-lived RAB-3 marked pre-SVs are observed to be stationary across a 10μm stretch of the TRN process (Figure 1F). This density is independent of both GFP::RAB-3 concentration and the presence of endogenous RAB-3 (Figure S1H). Motor neuron commissures in *C. elegans* have a lower density of long-lived stationary pre-SVs compared to that in TRNs (Figure 1F), while the density of RAB-5 marked long-lived stationary endosomal compartments is ~2±1.0 across 10μm of the TRN process (Figure 1F).

**Figure 1:**
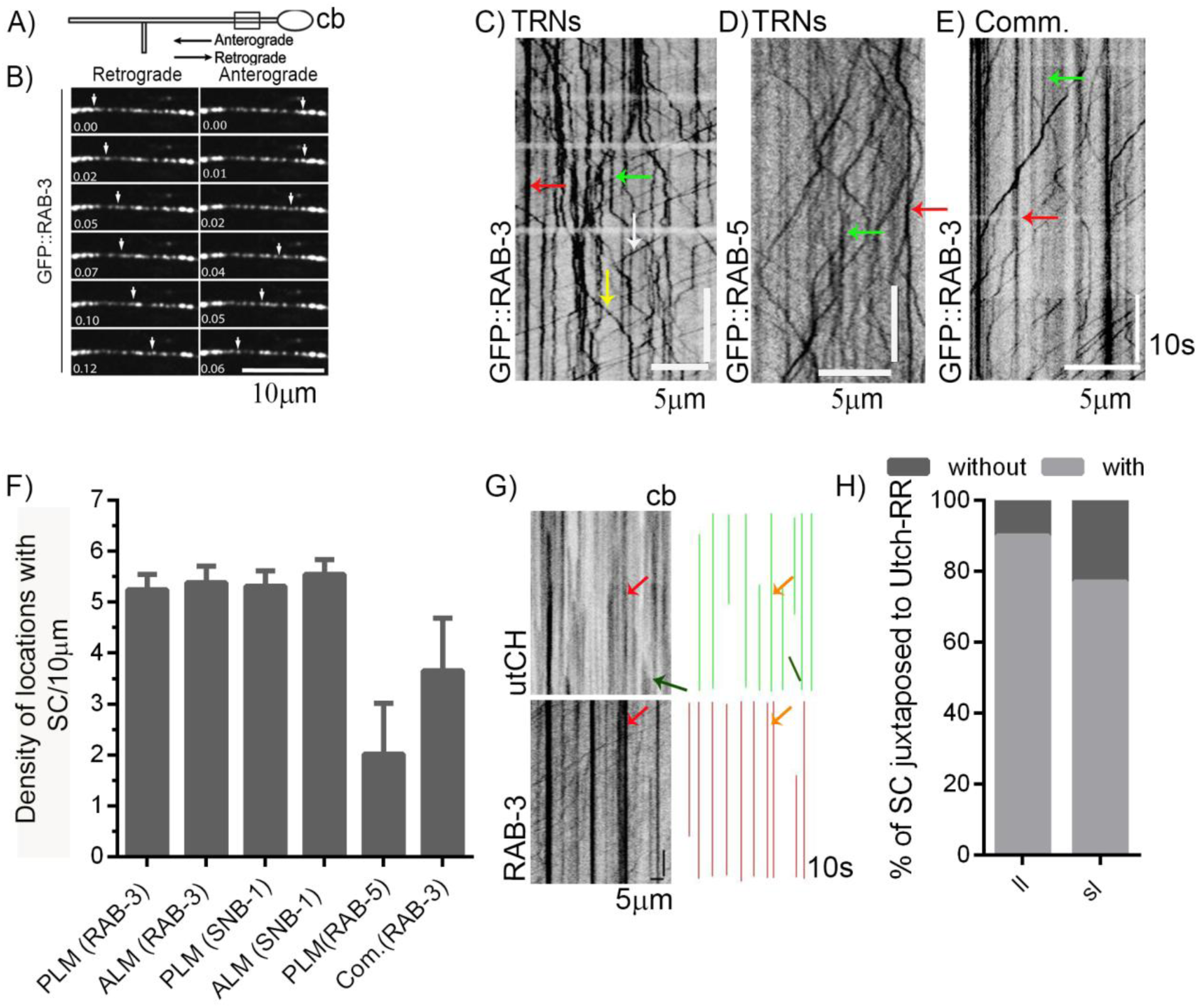
Stationary cargo are juxtaposed to immobile F-actin enriched regions. A) Schematic representation marking the region imaged in TRNs of *C. elegans* (*C.e*) B) Time lapse images showing retrograde (left) and anterograde motion (right) of GFP::RAB-3 tagged pre-SVs (white arrow) in a *C.e* TRN Kymograph representation of time-lapse movies of: C) pre-SVs tagged with GFP::RAB-3 in a *C.e* TRN, D) endosomal cargo tagged with eGFP::RAB-5 in a *C.e* TRN, E) pre-SVs tagged with GFP::RAB-3 in a *C.e* motor neuron commissure; Arrows: green: short-lived stationary cargo (SC), red: long-lived SC, yellow: retrograde cargo and white: anterograde cargo. Cell body (cb) is on the right hand side in all the kymographs F) Number of stationary vesicular cargo per 10μm observed in 75s time lapse movies of different cargo types and neurons of *C.e* (n=10 animals, number of 10μm bins analyzed ≥15, data represented as mean±SEM) G) Representative kymographs showing juxtaposition between GFP::utCH and mCherry::RAB-3 in *C.e* TRNs. Right panel shows the annotations of respective markers and yellow/red arrows represent an example where the two markers overlap. Green arrow in the right panel and green line in the left panel represent an actin trail. Cell body (cb) is on the right hand side in the kymograph. Note only long-lived stationary cargo and high intensity F-actin marked regions are annotated in the graphs. Data was acquired using simultaneous dual color time lapse imaging H) Percentage of stationary pre-SVs marked by RAB-3 juxtaposed to immobile GFP::utCH-rich regions (utCH-RR) in *C. e* TRNs (n≥12 animals for each, n=375 utCH-RR, n=228 stationary RAB-3 pre-SVs), ll: long-lived stationary cargo and sl: short-lived stationary cargo

In summary, we observe that various types of vesicular cargo can be stationary for several minutes and that the density of stationary cargo is specific to neuron and cargo type.

### RAB-3 marked vesicles accumulate and remain immobile for long durations at actin-rich regions

In cultured hippocampal neurons, both cortical actin and deep actin are known to be present throughout the neuronal process (Xu, Zhong et al. 2013, Ganguly, Tang et al. 2015). Stationary endosomes have been observed in these regions (Ganguly, Tang et al. 2015). Heterogeneities in the actin cytoskeleton could influence steady cargo flow. We therefore examined whether stationary pre-SVs along the neuronal process were present in actin-enriched regions. To test this, we examined the juxtaposition of stationary RAB-3 marked vesicles with actin-rich regions in *C. elegans* TRNs.

We used a transgenic line that expresses the GFP tagged calponin homology domain of F-actin binding protein Utrophin (utCH) as a marker of F-actin rich regions (Burkel, von Dassow et al. 2007, Chia, Patel et al. 2012, Chia, Chen et al. 2014). We observed mobile pools of F-actin, manifest as trails in our kymographs (Figure S2A, red arrow), in addition to immobile pools observed as high intensity vertical lines (Figure S2A, green arrow). These observations recapitulate ones made by Ganguly *et al* in cultured hippocampal neurons (Ganguly, Tang et al. 2015). The frequency of actin trails is 5±0.42 per 100μm per minute and their velocity of extension is 0.42±0.02μm/s. We also used Coronin-1::mCherry (COR-1), an orthologue of a mammalian actin binding protein associated predominantly with F-actin that is also known to cross-link actin and microtubules in yeast (Cai, Makhov et al. 2007, Wang, Zhou et al. 2013, Chen et al 2017). About half of the utCH rich regions are enriched with COR-1 as well (Figure S2B, C). This suggests that utCH and COR-1 mark an overlapping subset of actin-rich regions along the TRN process. COR-1 is less abundant along the neuronal process as compared to utCH and only 9% of the COR-1 enriched regions do not overlap with utCH (Figure S2C).

We tested the juxtaposition of stationary vesicular cargo with actin rich regions using dual color time-lapse imaging experiments. 90% and 77% of RAB-3 marked long-lived and short-lived stationary pre-SVs respectively are associated with utCH-rich regions (Figure 1G, H). The Manders Overlap coefficient value (see methods) measuring the spatial overlap between RAB-3 and Utrophin has a mean of 0.84±0.05 (Mean±SD). The values of the Manders Correlation coefficient, represented as (mean, standard deviation) are: M1 = (0.84, 0.04) and M2 = (0.86, 0.01). Analysis of the distribution of correlations yields evidence for non-zero means and positive skewness with a significance value of p < 0.05 for all 7 worms analyzed. These analyses suggest that the overlap between the two markers is significant and does not occur merely by chance. Nearly 45% of long-lived and 33% of short-lived stationary pre-SVs are juxtaposed to COR-1 enriched regions (Figure S2D, E). This difference in extent of juxtaposition of stationary pre-SVs *vis a vis* the two actin-markers likely arises due to a difference in their relative abundances along the neuronal process. The density of long-lived stationary RAB-3 marked cargo in a region depends linearly on the density of immobile utCH or COR-1 in the same region (Figure S2F, G). Although 80% of all stationary pre-SVs are associated with utCH marked actin-rich regions, only 59% of utCH marked regions associate with RAB-3 marked stationary pre-SVs. 73% of COR-1 rich regions associate with stationary pre-SVs. Thus, there are several actin-rich regions that do not have stationary pre-SVs associated with them.

It has been proposed that actin is nucleated from endosomes along the neuronal process (Ganguly et al 2015). We therefore examined the locations where actin polymerizes, observed as utCH trails in our kymographs (Figure S2A, red arrow). There is no significant bias in the actin trails where about 42% of the utCH trails occur at actin-rich regions lacking pre-SVs and about 47% of the utCH trails occur at regions where both stalled vesicles and actin are present (Figure S3A). Only 6% of all the actin polymerizing events occur at stationary RAB-3 vesicles not associated with actin (Figure S3A). We also observe that the utCH signal both appears and disappears at multiple locations along the TRN process (Figure S2A, cyan and magenta arrows). Only 23% of the appearance and disappearance events of utCH signal occur in proximity to stationary RAB-3 marked vesicles not initially associated with actin (Figure S3A). A majority of actin trails and appearance/disappearance of actin occur in regions without any stationary RAB-3 vesicles (Figure S3A). In a few stationary RAB-3 marked vesicles not associated with actin, we observe that utCH enrichment can appear transiently and then disappear with no effects on the location or intensity of stalled pre-SVs. Thus stalled vesicles may not be a major source of actin enrichment along the neuronal process.

In summary, nearly 90% of the long-lived stationary pre-SVs are present at utCH marked F-actin enriched regions. Only about 10% (20%) of long-lived (short-lived) stationary pre-SVs respectively are not associated with actin-rich regions. These stationary pre-SVs may be present at the ends of microtubules, which have been previously reported to lead to short-term cargo pausing *in vivo* (Yogev, Cooper et al. 2016) in other *C. elegans* neurons, or may be associated with the few COR-1 enriched sites lacking utCH.

### Multiple types of cargo accumulate at actin-rich regions

To examine whether multiple cargo types are associated with actin-rich regions, we investigated two additional cargo, RAB-5 and mitochondria in *C. elegans* TRNs. We observe that ~91% of immobile mitochondria and ~63% of immobile RAB-5 marked endosomes are associated respectively with utCH-rich and COR-1-rich regions (Figure 2A-C). Moreover, 95% of stationary mitochondria and 56% of stationary RAB-5 marked endosomes respectively also juxtapose with RAB-3 marked stationary pre-SVs (Figure 2D-F). The Manders Overlap Coefficient measuring the spatial overlap between RAB-5 marked stationary endosomes and RAB-3 marked stationary pre-SVs is 0.82 (SD 0.09). In addition, the values of the Manders Correlation coefficient, represented as (mean, standard deviation) are: M1 = (0. 84, 0.04) and M2 = (0.81,0.01). Analysis of the distribution of correlations yields evidence for non-zero means in 8 of the 10 worms studied at the p < 0.05 level, and positive skewness with a significance value of p < 0.05 for all 10 worms studied. These analyses suggest that the overlap between the two markers is significant despite their difference in abundance and does not occur by chance. Our data suggest that multiple types of cargo stall at the same regions.

**Figure 2:**
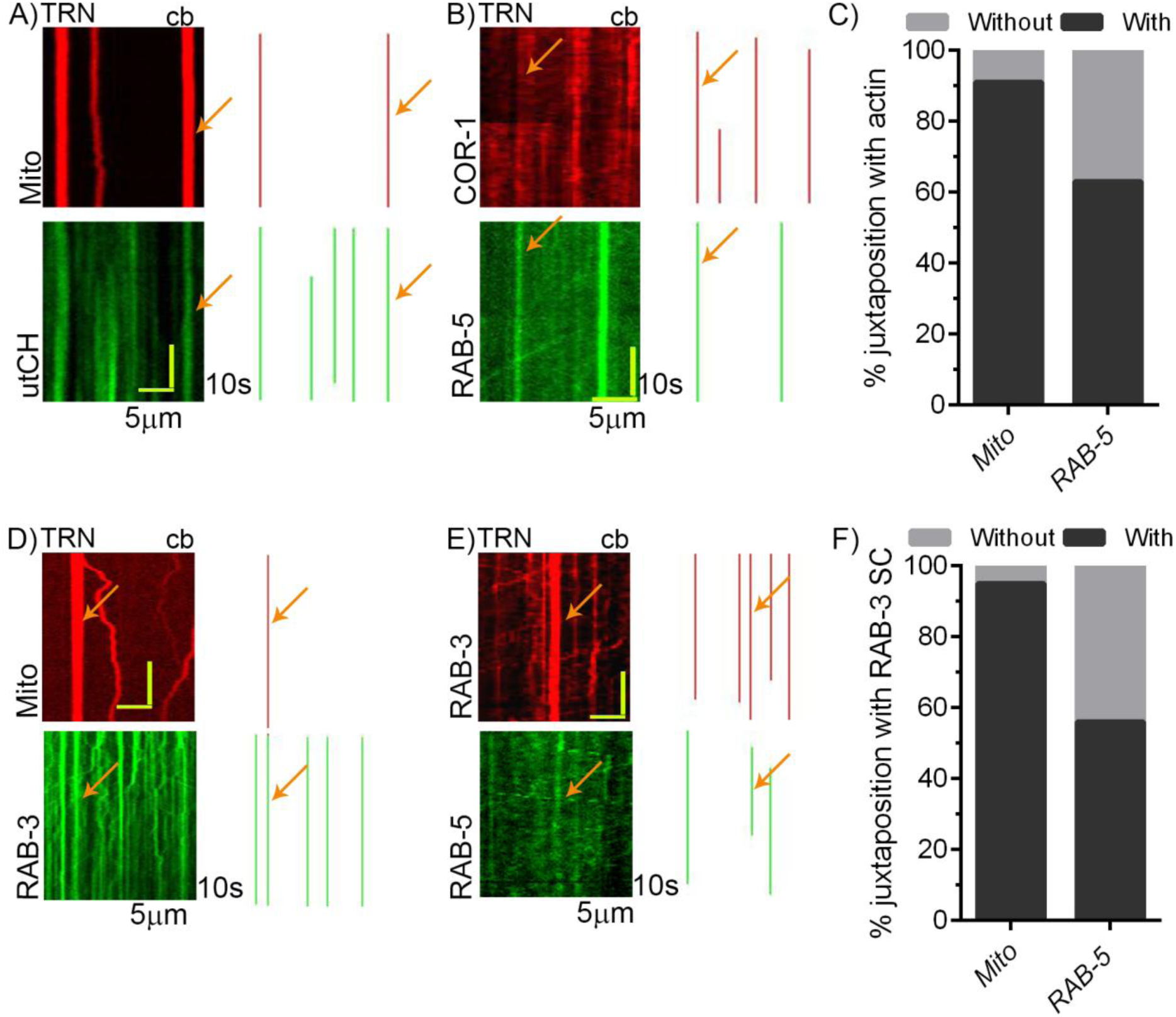
Multiple types of cargo are present at the same locations along the *C. elegans* TRN neuronal process. Representative kymographs (left) and their respective annotations (right) showing juxtaposition between A) tagRFP::MLS, mitochondria associated protein (Mito) and utCH::GFP, B) COR-1::mCherry and eGFP::RAB-5. The Manders overlap coefficient, represented as (mean, standard deviation), is (0.96, 0.03) for COR-1::mCherry and eGFP::RAB-5. The Manders correlation coefficients M1 and M2 are (0.72, 0.01) and (0.72, 0.02) respectively. Analysis of the distribution of correlations yields evidence for non-zero means and positive skewness with a significance value of p < 0.05 for all worms studied. Orange arrows represent an example where the two markers overlap. Data was acquired using simultaneous dual color time lapse imaging C) Percentage of mitochondria (Mito) and eGFP::RAB-5 marked endosomes juxtaposed to immobile actin-rich regions (Actin-RR) (n=10 animals, n=58 mitochondria, n=234 utCH rich regions, n= 95 RAB-5 marked vesicles, n= 173 COR-1 enriched regions) Representative kymographs (left) and their respective annotations (right) showing juxtaposition between D) GFP::RAB-3 and tagRFP::MLS marked mitochondria (Mito), E) RAB-3::mCherry and eGFP::RAB-5. Orange arrows represent an example where the two markers overlap F) Percentage of mitochondria (Mito) and eGFP::RAB-5 marked endosomes present with and without RAB-3 marked stationary pre-SVs (n=10 animals, n=70 mitochondria, n=200 RAB-3 marked vesicles; n= 145 RAB-5 marked vesicles, n=178 RAB-3 marked vesicles) Note: in all the overlay images representing juxtaposition between two markers, only long-lived stationary cargo and high intensity of the F-actin marker signals are annotated. In all the kymographs, cell body (cb) is on the right hand side. Data was acquired using either simultaneous or sequential dual color time lapse imaging

We also attempted three color imaging to directly observe two different types of cargo stalled at a location enriched with actin. However, the use of three markers whose expression was driven in the same cell resulted in a nearly 99% reduction in flux of all tagged cargo-types along the neuronal process. This made quantitation and interpretation of results difficult and hence we did not attempt any further three-color imaging experiments. We also observe a reduction in flux in dual color imaging compared to imaging the same transgene as a single marker (Figure S3B). However, the density of SC in animals expressing two transgenes remains unchanged (Figure S3C).

Since multiple cargo, RAB-3 and RAB-5 marked vesicles as well as mitochondria all halt at the same actin-rich locations, we infer that cargo experience common constraints to their motion in a complex environment.

### Disrupting the actin cytoskeleton reduces the density of stationary RAB-3 marked vesicles

To test the contribution of the actin cytoskeleton in accumulation of stationary vesicular cargo, we injected Latrunculin A (LatA) into the body cavity of *C. elegans* expressing GFP::RAB-3 in their TRNs. We observe that in wild type animals, the length of each utCH marked region extends typically to ~1.2μm±0.8μm along the neuronal process, with ~46% spanning a 0.6 to 1.4μm region (Figure S3D, 4A). On LatA injection, utCH-rich regions become narrower and the number of utCH enriched regions spanning 0.6 to 1.4μm drops to 28% (Figure S3D, 4A). We also observe that the density of immobile COR-1 and immobile utCH marked F-actin enriched regions reduces by 30% without significant changes in flux, concomitant with a corresponding ~30% reduction in the density of long-lived stationary pre-SVs (Figure 3C, Figure S4B, C, 5A).

**Figure 3:**
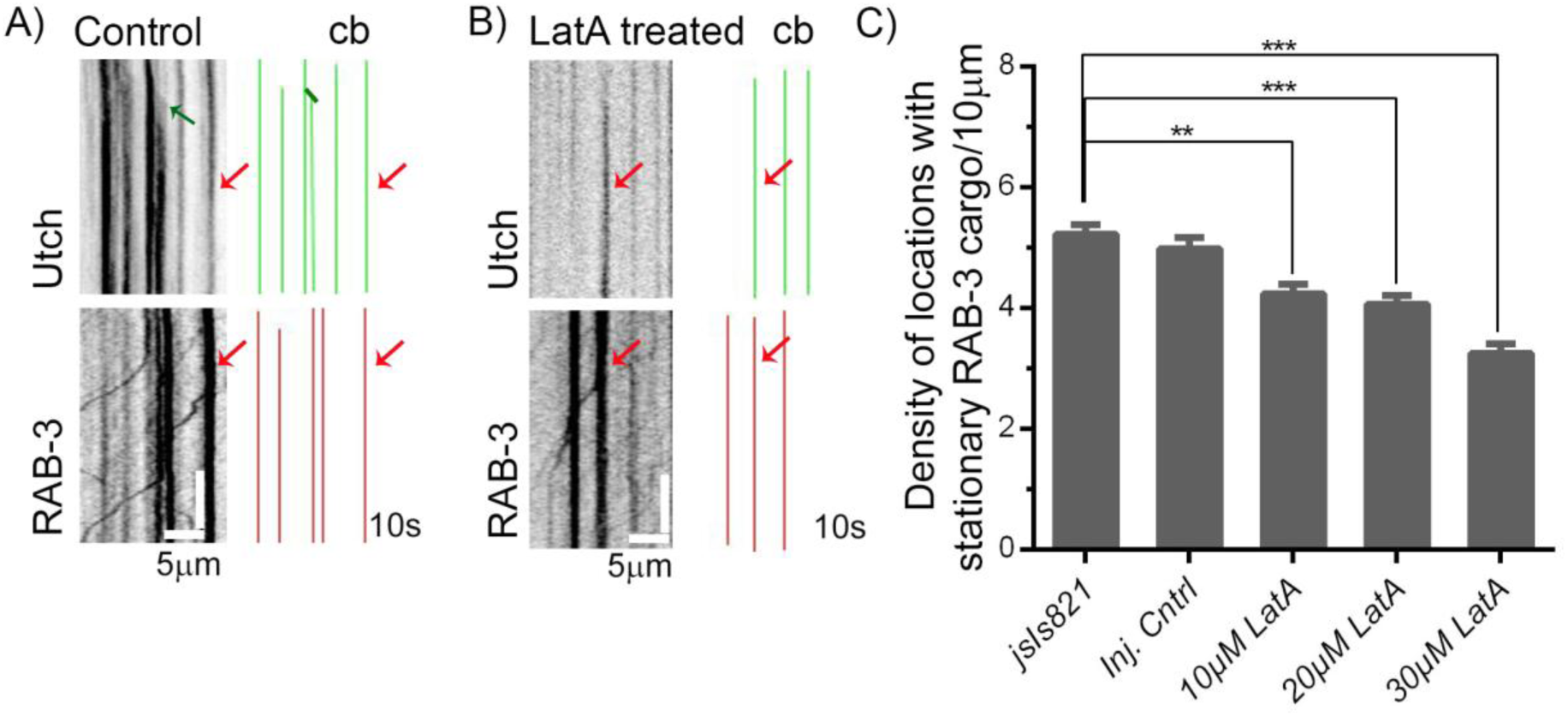
Perturbation of actin-rich regions reduces the density of stationary pre-SVs in *C. elegans* TRNs. Representative kymographs of utCH::GFP and RAB-3::mCherry from A) control and B) Latrunculin A (LatA) treated animals. Note: same criterion as mentioned in Figure 1 and 2 was used to annotate overlay images representing juxtaposition between two markers. Red arrows represent an example where the two markers overlap. Both green arrow and green line in A) represent actin trails. Cell body (cb) is on the right hand side in kymographs. The juxtaposition between utCH::GFP and mCherry::RAB-3 is significant, p<0.05, for all the LatA treated animals. Mander’s coefficient, Mean±SD is 0.91±0.051, M1 and M2 are (0.75, 0.02) and (0.74, 0.02) respectively. Data was acquired using simultaneous dual color time lapse imaging. A larger region of the neuron showing actin dynamics in control and stimulated animals is shown in Fig. S3D) C) LatA concentration dependant reduction in the density of GFP::RAB-3 stationary cargo. (n≥ 7 animals for each concentration, data represented as mean±SEM, One way ANOVA with Dunn’s multiple comparison test was used for comparison. All the values were compared with buffer injected controls, * * p<0.01, * * * p<0.001)

The extent of reduction in density of stationary pre-SVs is directly proportional to the concentration of LatA injected (Figure 3C). Nearly half of the actin-rich regions that persist after LatA treatment continue to have stationary RAB-3 marked vesicles associated with them. On the other hand, the percentage of stationary RAB-3 marked pre-SVs that are not associated with actin-rich regions almost doubles (Figure S4D). This may reflect actin-independent sources of stationary vesicles that persist after LatA treatment, for example at microtubule ends (Yogev, Cooper et al. 2016).

We examined whether the reduction in density of stalled pre-SVs in LatA treated animals contributes to the mobile pools of vesicles. We first compared the overall flux of GFP::RAB-3 marked vesicles in both injection controls and LatA treated animals. Buffer injected controls have slightly reduced flux when compared to uninjected controls (Figure S5A). LatA injected animals also have slightly reduced flux compared to injected controls (Figure S5A). However, we cannot assign statistical significance to these changes, since the process of injection itself leads to significant variability in flux (Figure S5A). We therefore instead determined the fraction of vesicles that are moving or stationary for each treatment. The number of stationary vesicles in each kymograph was determined by calculating the intensities of regions with stationary vesicles divided by the average intensity of individual paused vesicles in the same analyzed kymograph. Control animals have about 30% of vesicles that are stationary and the remaining 70% are moving (Figure S5B). Across LatA treatments the percentage of stationary vesicles vary from 12%-22% and the motile vesicles vary between 78%-88% (Figure S5B). These differences although small suggest that when actin is depolymerized, the numbers of stationary vesicles reduce. These formerly stationary vesicles may contribute to the motile fraction.

Our data suggest that accumulation of stationary pre-SVs along the neuronal process depends on the presence of actin-rich regions and that the density of stalled pre-SVs reduces upon actin disruption. The reduced density of stalled pre-SVs after actin depolymerization may contribute to pools of moving vesicles.

### Moving vesicles stall more frequently at actin-rich regions associated with stationary cargo

We showed that vesicular cargo are stationary for long periods at regions enriched in F-actin. In the narrow geometry of a *C. elegans* neuron (Chalfie and Thomson 1982, Cueva, Mulholland et al. 2007), such actin-rich locations alone along with actin-rich locations associated with stalled cargo can both impede the movement of cargo. To test this hypothesis, we compared the behavior of moving pre-SVs when they encounter actin-rich regions with and without pre-existing stalled vesicles.

We analyzed dual color movies with F-actin enriched regions marked by utCH::GFP and RAB3-mCherry marked pre-SVs. At stable F-actin enriched locations that are associated with stalled RAB-3 cargo, 79% of the anterogradely and retrogradely moving pre-SVs stop. 17% of moving pre-SVs stop on encountering a location enriched with F-actin alone while only 5% of RAB-3 marked vesicles stop at a location that has neither F-actin nor a pre-existing stationary pre-SVs (Figure 4A). 14% of the transient utCH appearance events lead to new stationary cargo forming at the region after utCH appears along the process (Figure S2A, S5C). This suggests that actin-rich regions along the neuronal process likely act as hotspots where pre-SVs tend to stall. Vesicles stalled at actin-rich regions appear to act as more effective roadblocks for moving vesicles, stalling them at higher frequencies than in actin-rich regions alone. Stalled RAB-3 pre-SVs alone in the absence of actin are also able to cause substantial stalling of other RAB-3 marked pre-SVs although less than in regions that contain actin (compare 80% to 60% in Figures 4A and 4B). We find no changes in the number of vesicles that stop at locations with stationary vesicular cargo across several developmental stages. (Figure S5E, F).

**Figure 4:**
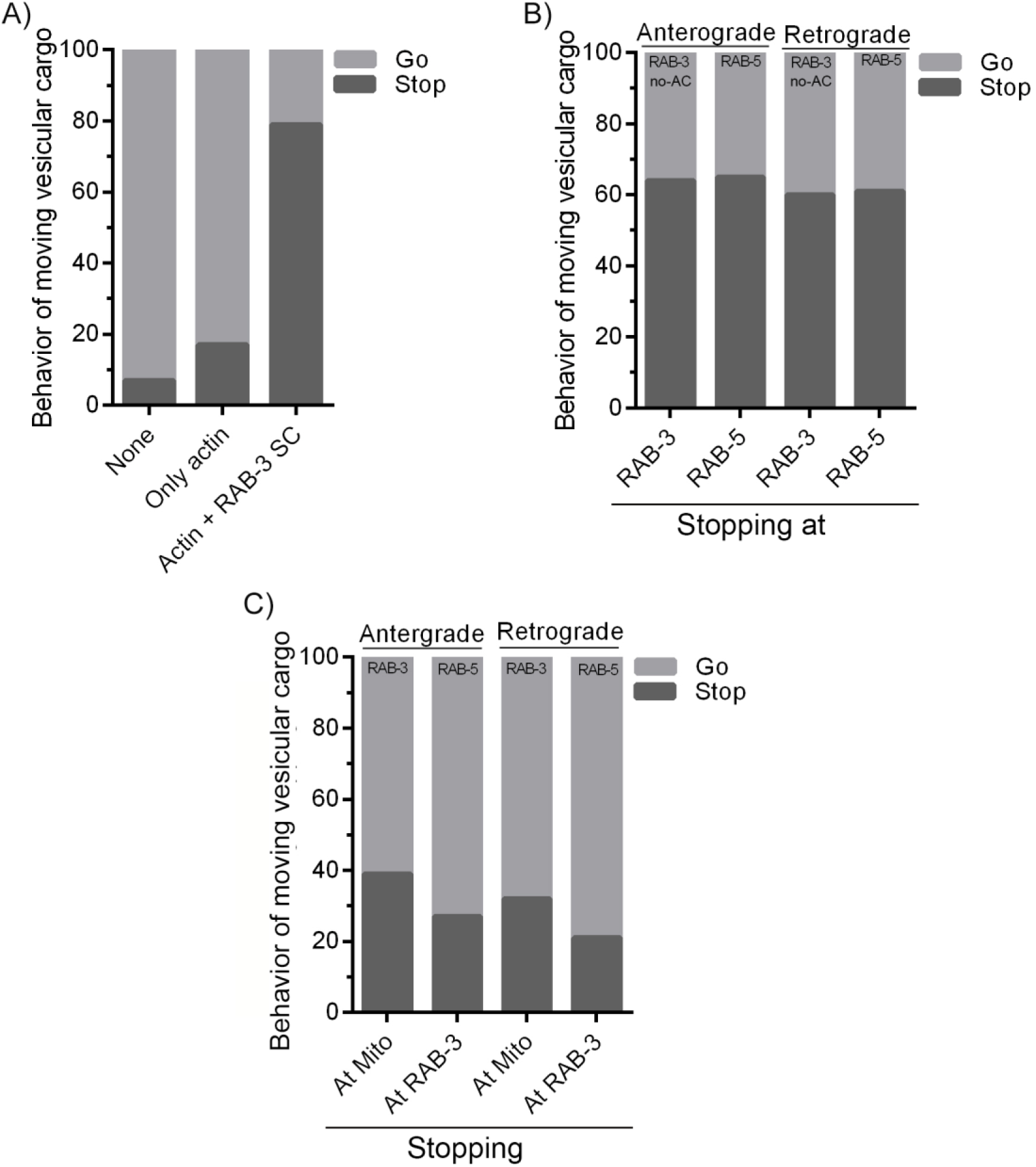
Stationary RAB-3 pre-SVs at actin-rich regions act as local roadblocks to moving RAB-3 pre-SVs in *C. e* TRNs. A) Percentage of cargo stopping at or continuing to move through regions that are neither actin-rich nor associated with pre-existing stationary vesicles, regions that are only actin-rich, and actin-rich regions with a stationary pre-SV (SC). (Increase in stopping at actin-rich regions associated with stationary cargo compared to regions enriched only with actin is statistically significant, One way ANOVA followed by multiple comparison, n=10 animals, number of vesicles analyzed ≥100, *p<0.05. See materials and methods for the details of analysis) B) Percentage of cargo stopping at locations occupied by cognate stationary cargo in both anterograde and retrograde direction. The bar graphs represent percentage of moving RAB-3 marked pre-SVs stopping at stationary RAB-3 marked pre-SVs lacking actin (no-AC) and percentage of moving RAB-5 endosomes stalling at RAB-5 stationary vesicles lacking immobile RAB-3 marked pre-SVs. (n=10 animals each, number of vesicles analyzed ≥ 40) C) Percentage of cargo stopping at locations occupied by non-cognate stationary cargo in both anterograde and retrograde directions. The bar graphs represent percentage of moving RAB-3 marked pre-SVs stopping at stationary mitochondria (Mito) lacking stalled RAB-3 marked pre-SVs and percentage of moving RAB-5 endosomes stalling at RAB-3 stationary pre-SVs lacking immobile RAB-5 marked endosomes. (n=12 animals each, number of vesicles analyzed ≥ 40)

Since we observe a reduction in flux of pre-SVs in animals expressing two transgenes, we examined if the nature and fraction of events observed at stationary cargo are influenced by reduced flux. We see that a comparable fraction of mCherry::RAB-3 marked vesicles stop when they encounter a location with stalled pre-SVs in animals expressing either the mCherry::RAB-3 transgene alone or both mCherry::RAB-3 and utCH transgenes (Figure S5D).

We examined the time taken for a RAB-3 marked vesicle that stops at a location with pre-existing stationary RAB-3 vesicles to begin moving again. We photobleached stationary cargo marked by GFP::RAB-3 to clearly visualize moving cargo stalling at these locations. Out of 52 vesicles tracked, 45 vesicles stall and do not begin moving again. The remaining 7 are observed to move again after pausing for 3±2s. This average pause duration at regions with stalled vesicles is greater than the 1.4±0.54s average pause duration of RAB-3 marked vesicles in regions lacking stationary RAB-3 vesicles (Table S2).

We further investigated whether the stopping of moving vesicles in regions where other vesicles were stalled depended on the type of stationary cargo such moving vesicles encountered. We use dual color time-lapse imaging to label two different cargo types. In each case, we assessed stopping at regions lacking their cognate stationary cargo. In *C. elegans* TRNs, we imaged RAB-3 marked pre-SVs and mitochondria, RAB-3 marked pre-SVs and RAB-5 marked endosomes. We observe that ~6% of mitochondria have no associated stationary pre-SVs at the beginning of the movie (Figure 2F). We tracked every moving RAB-3 marked vesicle encountering a mitochondrion with no associated stationary pre-SVs. We observe that 39% and 27% of the anterogradely and retrogradely moving vesicles respectively, stop after encountering an immobile mitochondria (Figure 4C). We similarly observe that 32% of anterogradely moving and 20% of retrogradely moving RAB-5 marked endosomes stall at sites occupied by stationary RAB-3 marked pre-SVs (Figure 4C). Such stalling of moving vesicles at non-cognate sites suggests that immobile cargo of any type along the neuronal process can act to stall movement of any other cargo. A greater proportion of RAB-5 (60%) marked vesicles stall at locations with stationary RAB-5 vesicles that lack RAB-3 stationary cargo (Figure 4B). These data suggest that there might be additional cargo specific mechanisms leading to greater cargo accretion.

In summary, actin-rich regions, actin rich-regions with stalled cargo and stalled cargo all appear to stop cargo moving through a given region of the neuronal process causing local traffic jams. Our data suggest that cargo crowding especially at sites enriched with actin play a key role in local traffic jams along the neuronal process.

### The presence of stationary cargo locally impedes cargo motion

Our data show that moving cargo stall more frequently at actin-rich regions with associated stalled cargo and at stalled cargo, suggesting that the presence of stalled cargo can impede cargo movement. We thus examine: 1) whether the majority of cargo stalling occurs where stationary cargo pre-exist, 2) whether mobilization of stalled vesicles leads to local increases in local cargo flux and 3) if the density of stalled cargo in a region correlates with run-lengths of moving cargo.

Using GFP::RAB-3 and SNB-1::GFP movies in TRNs and GFP::RAB-3 in motor neuron commissures, we observe that only 5% of pre-SVs stall away from pre-existing stationary pre-SVs in both neuron types (Figure 5A). To address the mobilization of stalled vesicles we specifically examine locations where stationary cargo mobilize and disperse in our 5 minute time lapse movies of GFP::RAB-3 (Figure 5B). We count the number of moving pre-SVs that stop or continue moving through the same location before and after stationary pre-SVs mobilize. As expected, there is a substantial increase of local cargo flux at the location after stationary pre-SVs mobilize (Figure 5B, C). This increase is approximately three-fold with no appreciable change in the number of vesicles that encounter the region before or after the stationary pre-SVs mobilize (Table S3). Finally, to assess whether the density of stalled cargo in a region correlates with run-lengths of moving cargo, we chose non-overlapping 20μm regions on kymographs with differing densities of stationary pre-SVs, calculating the distance a given cargo in these regions moves before stopping. We find that total run lengths of moving pre-SVs in a region are inversely proportional to the density of stationary pre-SVs (Figure 5D).

**Figure 5:**
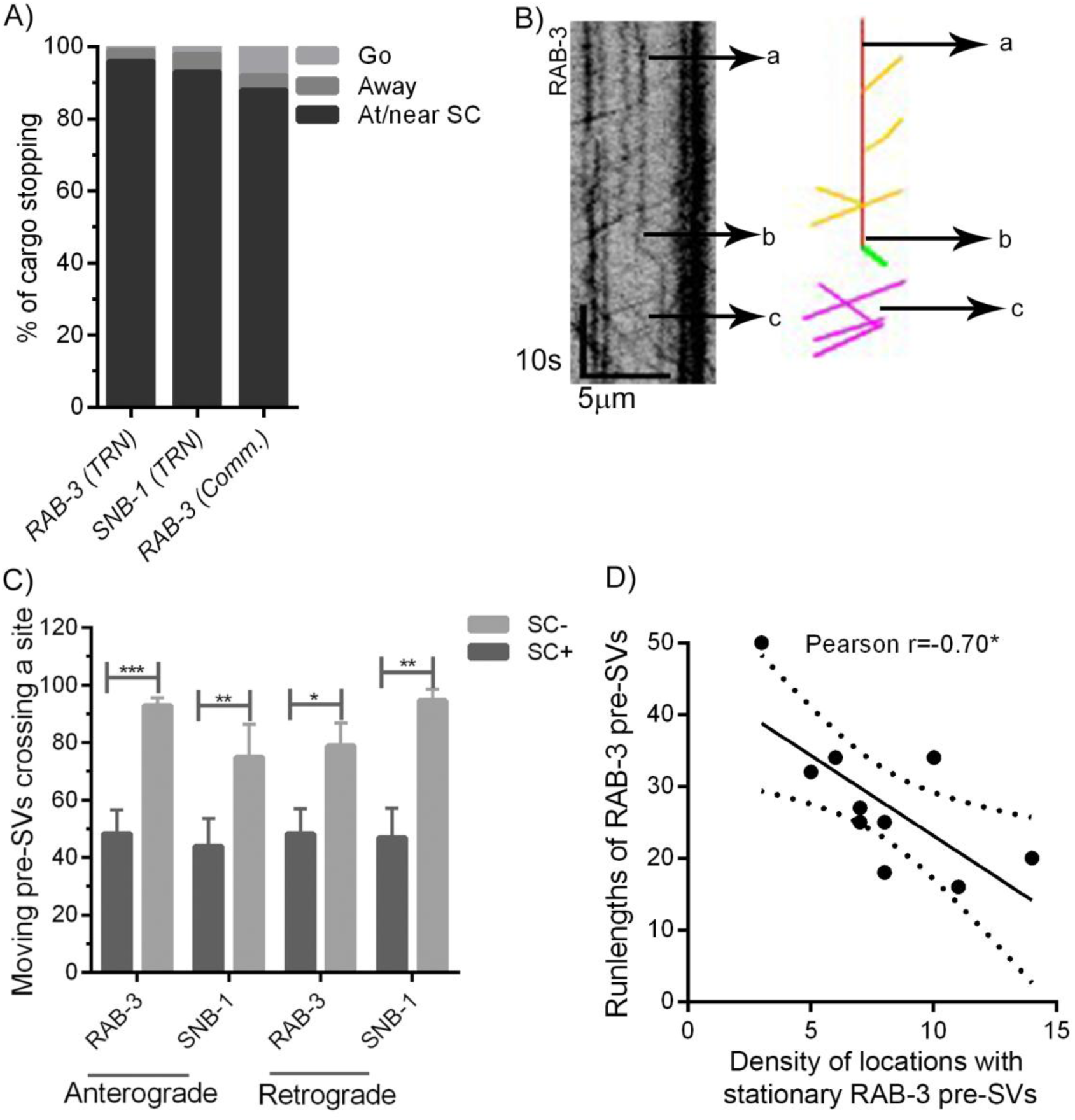
Stationary cargo locally modulate motion of moving cargo. A) Percentage of GFP::RAB-3 marked cargo stopping away from stationary RAB-3 cargo (SC) in the neuronal process of *C.e* TRNs and *C.e* motor neurons commissures (Comm.) (n=10 animals, number of vesicles analyzed ≥200) B) Kymograph illustrating the quantitation of flux at a location from where a stationary RAB-3 pre-SV has mobilized in *C.e* TRN; a) when stationary pre-SV was present, b) stationary pre-SV mobilized, and c) after stationary pre-SV mobilization. Overlay image of events (right) occurring is shown for clear visualization. Red line: stationary pre-SV, green line: mobilization of stationary pre-SV, yellow and magenta line: trajectories of moving vesicles before and after mobilization of stationary cargo respectively C) Percentage of moving vesicles crossing a site in the presence of a stationary cargo (SC+) and after it has mobilized (SC-) in *C.e* TRN ( paired t-test, n=10 animals, number of vesicles analyzed ≥50, number of sites=20, Data represented as Mean±SEM) D) Regression plot between the density of stationary pre-SVs and average run-length of moving pre-SVs marked by GFP::RAB-3 in the same region of *C.e* TRN. Dotted lines in the two graphs represent 95% confidence band for the best-fit line (n=10 animals analyzed, number of regions=14, n=80 vesicles, *p<0.05)

To summarize, stalling of vesicles occurs largely at locations with pre-existing stationary cargo suggesting that local obstruction of transport might occur preferentially at such locations. Local increases in flux after stationary pre-SVs mobilize, and the dependence of distance travelled on the density of stationary pre-SVs, are consistent with locally impeded transport where the locations with stalled cargo act as roadblocks for any other moving vesicle at that location.

### Cargo transport is impeded at actin-rich regions in *Drosophila* neurons

To test if actin-rich regions associated with stalled cargo result in local traffic jams of microtubule based cargo transport in another model system, we examined axons of *Drosophila* chordotonal neurons. We used dual color imaging to investigate: 1) the juxtaposition between stationary vesicular cargo and actin-rich regions, marked by RAB-4-mRFP (Salvaterra and Kitamoto 2001) and Lifeact-GFP (Riedl, Crevenna et al. 2008) and 2) the behavior of moving cargo through actin-rich regions with stalled RAB-4 marked vesicles.

We observe that about 70% of the long-lived stationary RAB-4 marked vesicles are found close to Lifeact-rich regions and that 70% of Lifeact enriched locations have stationary RAB-4 marked cargo next to them (Figure 6A-C). The Manders Overlap Coefficient between RAB-4 marked vesicles and Lifeact-enriched regions has a mean of 0.94 (SD 0.05). In addition, the values of the Manders Correlation coefficient, represented as (mean, standard deviation) are: M1 = (0.87, 0.04) and M2 = (0.86, 0.04). Analysis of the distribution of correlations yields evidence for non-zero and positive means in 8 of the 10 larvae imaged at p < 0.05, and positive skewness with a significance value of p < 0.05 for 7 of the 10 larvae imaged. These analyses suggest that the overlap between the two markers is significant. The density of long-lived stationary vesicular cargo in a region depends linearly on the number of Lifeact-rich locations in that region (Figure 6D). As observed in *C. elegans* TRNs, LatA treatment reduces the percentage of Lifeact-rich regions by 30% with a corresponding reduction in the density of stationary vesicles (Figure 6E, Figure S4C). This reduction occurs without a change in flux of RAB-4 tagged vesicles (Figure S5A). These data suggest that actin-rich regions contribute to the presence of stationary vesicular cargo in *Drosophila* chordotonal neurons as well.

**Figure 6:**
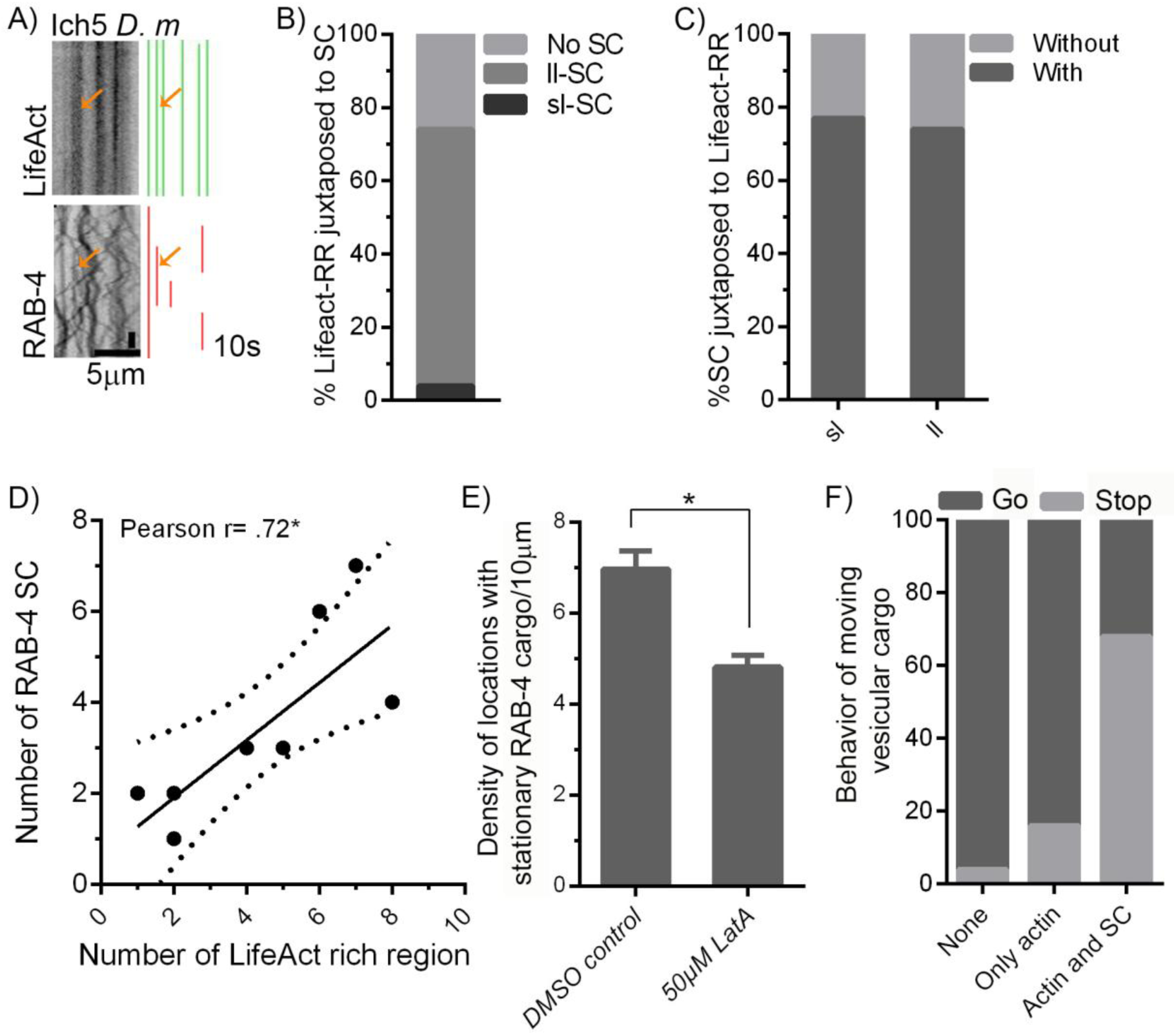
Regulation of moving cargo at F-actin enriched locations in chordotonal neurons of *D. melanogaster*. A) Left: kymographs made from movies of Lifeact-GFP and RAB-4-mRFP. Right: annotations of two markers used for juxtaposition analysis. Cell body is on the right hand side in kymographs. Note: same criterion as mentioned in Figure 1 and 2 was used to annotate overlay images representing juxtaposition between two markers. Orange arrows represent a location where both markers are present B) Percentage of Lifeact-rich regions (Lifeact-RR) associated with RAB-4 stationary vesicular cargo (SC), sl- short-lived, ll-long-lived (n=10 animals, n= 167; Lifeact-RR, n= 115 vesicles) C) Percentage of RAB-4 stationary cargo (SC) juxtaposed to Lifeact rich regions (Lifeact-RR) (n=10 animals, n= 160; Lifeact-RR, n= 196 vesicles) D) Regression plot of density of Lifeact enriched regions and RAB-4 stationary cargo (SC) (n=9animals, p<0.01). Dotted lines represent the 95% confidence band of the best-fit line, * p<0.05 E) Density of stationary vesicular cargo marked by RAB-4-mRFP per 10μm in control animals and animals treated with 50μM LatA (Data represented as Mean±SEM, n=10 animals and 10 bins analyzed, Student’s t-test, * p<0.05) F) Percentage of cargo stopping at or continuing motion without interruption after encountering heterogeneous locations along the neuronal process (Increase in stopping at actin-rich regions associated with stationary cargo compared to regions enriched only with actin is statistically significant, One way ANOVA followed by multiple comparison, n=10 animals, n=200 vesicles, * p<0.05. See materials and methods for the details of analysis) All data was acquired using sequential dual color time lapse imaging

To investigate whether actin-rich regions with stationary RAB-4 cargo act to stall moving RAB-4 cargo we carried out the same comparative analysis described earlier. We observe that on encountering a location enriched with both RAB-4 marked stalled cargo and Lifeact, ~70% of moving RAB-4 vesicles stop. Around ~15% of moving vesicles stop on encountering regions enriched with Lifeact but without RAB-4 marked cargo. As with TRNs and commissures in *C. elegans,* the majority of moving cargo either marked by RAB-4 or SYT-1 stall predominantly at SC in *Drosophila* neurons as well (Figure S6A). Since *C. elegans* TRNs exhibit similar obstruction of transport at regions that are both actin-rich and have pre-existing stalled vesicles (Figure 4A), impeded cargo transport at such locations can be argued to be a general phenomenon associated with axonal transport of cargo.

### Stationary vesicular cargo are dynamic reservoirs of vesicles

Our kymographs show that more than 95% of vesicles moving over a 70-80μm length of the PLM will eventually encounter pools of stationary vesicular cargo and stop (Figure 5A). Unless compensated by the remobilization of stationary cargo, this is likely to lead to a steady reduction in cargo flux along the neuronal process over time.

To determine if moving vesicles can arise from stationary vesicles, we first examined the numbers of vesicles present at such vesicular clusters. We find that there are approximately 2-4 vesicles in stationary vesicle clusters of RAB-3-marked pre-SVs in *C. elegans* TRNs as well as RAB-4-marked vesicles in *Drosophila* neurons, based on the distribution of intensities of individually paused vesicles (Figure 7A, Figure S6B). Consistent with this, we observe that when stationary vesicles disperse completely, they release 2-4 vesicles (Figure 7B). Similar small groups of synaptic vesicles have been observed in EM sections in asynaptic sites along the axons of HSN neurons of *C. elegans* (Shen, Fetter et al. 2004) and may be similar to the clusters of stationary pre-SVs seen in this study.

**Figure 7:**
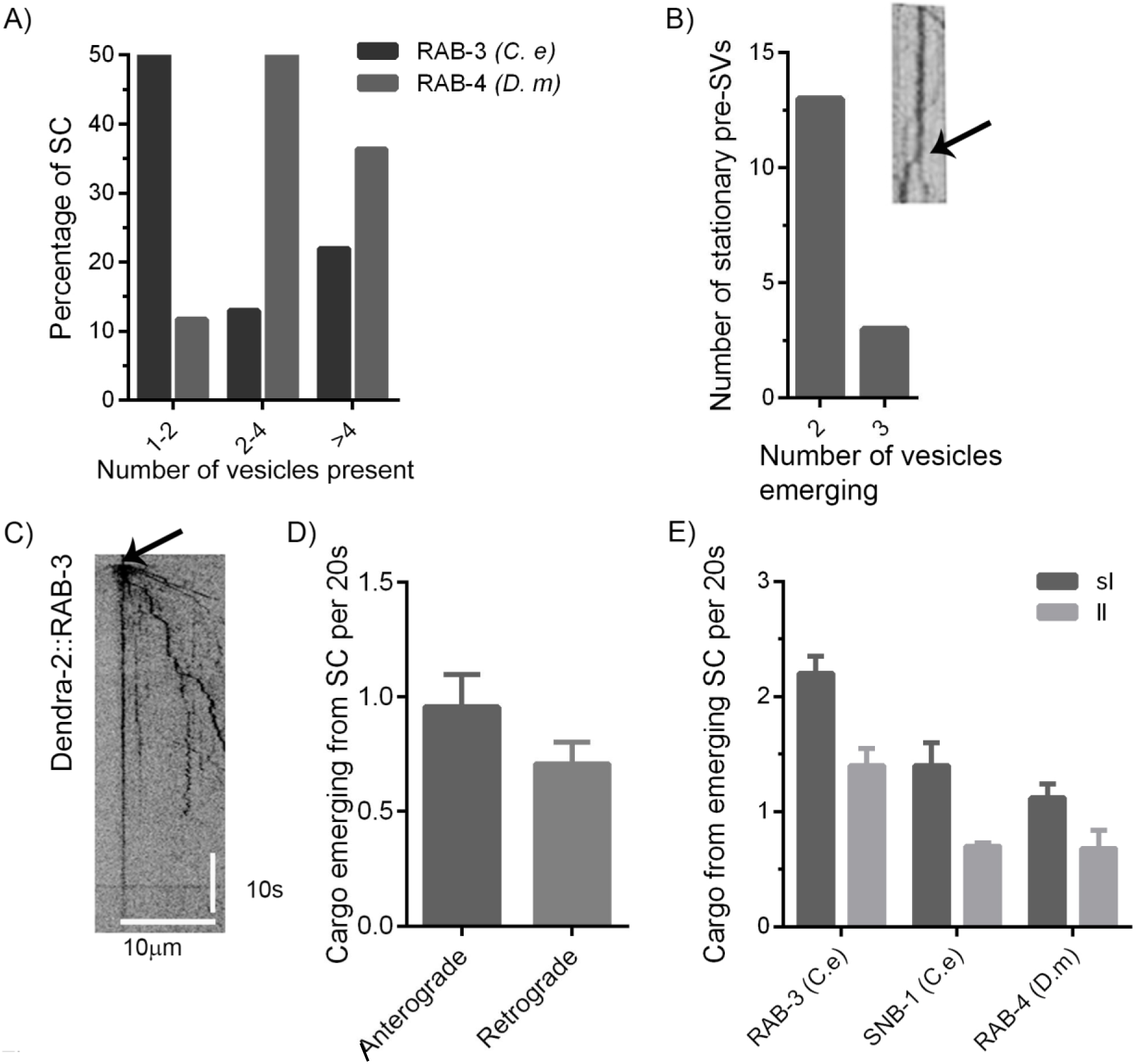
Stationary cargo can capture and release vesicles. Approximation of number of vesicles present at a stationary cargo cluster (SC) A) by counting number of vesicles that arise when a stationary cargo disperses into multiple vesicles in *C. e* TRN and B) by using average intensities of individual paused vesicles in both *C. elegans* TRNs and *D. melanogaster* chordotonal neurons (n ≥ 10animals, number of SC that mobilize≥ 15, number of SC≥20) C) Representative kymograph showing Dendra-2::RAB-3 photoconversion in *C.e* TRN D) Number of Dendra-2::RAB-3 marked vesicles emerging from a location containing stationary pre-SVs per 20 seconds after photoconversion, (n= 20 animals, number of stationary pre-SVs analyzed ≥30) E) Average number of vesicles emerging from a location containing stationary vesicular cargo (SC) in kymographs acquired from different neurons expressing vesicle markers (n ≥ 10 animals, number of stationary vesicular cargo analyzed ≥ 10)

Since several vesicles are present at stationary cargo clusters, we examined whether these locations contribute to flux along the neuron, using photoconversion of Dendra-2::RAB-3 (Figure 7C, D). We find that stationary pre-SV clusters release a vesicle approximately every 20 seconds (Figure 7D), a number comparable to that obtained from our conventional GFP::RAB-3 kymographs (Figure 7E). At a given location where RAB-3 marked vesicles are stationary, we observe that of the total anterogradely moving vesicles that encounter this location, 16% go past the location, 54% stop and about 46% vesicles emerge from the same pool of stalled vesicles (Figure S6F). More broadly, the ability of stationary clusters to hold and subsequently release vesicles appears to be a common feature of cargo transport in *Drosophila* neurons as well (Figure 7E).

Since the motion of pre-SVs depends on the UNC-104/KIF1A motor (Hall and Hedgecock 1991), we examined whether a change in levels of motor on the cargo surface resulted in altered pre-SV behavior at locations where cargo stall. We examined wild type animals, an *unc-104* mutant encoding Kinesin-3 motor defective in cargo binding with known reduction in levels of motor on the pre-SV surface (Kumar, Choudhary et al. 2010), and animals over-expressing the UNC-104 motor. A reduction in UNC-104/KIF1A does not change the locations where pre-SVs stall with ~77% of all stationary pre-SVs present at stable actin-rich regions, similar to wild type (Figure S6C). Further, the mutants do not show any change in the density of stable utCH tagged F-actin when compared to wild type (Figure S6D). In *unc-104* animals, the fraction of anterogradely moving RAB-3 marked vesicles that stop at locations with pre-existing stationary pre-SVs significantly increases (Figure S6F). In addition, the number of vesicles that emerge to move in the anterograde direction from a pre-existing pool of stationary RAB-3 vesicles is 9% lower than in wild type (Figure S6F). Retrogradely moving vesicles, as expected, do not show any changes in the fraction stopping (Figure S6G). Vesicles in animals over-expressing UNC-104 behave similar to wild type suggesting that sufficient motors may already be present on the cargo surface in wild type (Figure S6F, G).

We also attempted to address whether the stalled RAB-3 vesicles are associated with motors. We observe that 77% of locations with stationary RAB-3 pre-SVs show UNC-104::GFP enrichment (Figure S6E). These motors may be bound to pre-SVs, although it is difficult to exclude the presence of independent motor reservoirs associated, for instance, with actin at such locations. Since only 42% of moving vesicles co-migrate with UNC-104::GFP in our movies, it is difficult to assess whether UNC-104 is recruited on pre-SVs from regions that contain stationary RAB-3 vesicles (Figure S6E).

The ability of moving vesicles to incorporate with as well as emanate from the locations of stationary cargo suggests that these locations serve a natural function as dynamic reservoirs of vesicles along the entire neuronal process. Levels of motors on the cargo surface appear to play a role in both stalling as well as in mobilization of vesicles from stationary cargo clusters.

## DISCUSSION

Crowded environments pose special problems for steady transport. The axon is narrow and filled with cytoskeletal elements, associated proteins and other cargo, leading to a congested transport path that cargo must navigate (Chalfie and Thomson 1979, Chalfie and Thomson 1982, Hirokawa 1982, Black and Kurdyla 1983, Cueva, Mulholland et al. 2007, Kaasik, Safiulina et al. 2007, Jaworski, Hoogenraad et al. 2008, Leterrier, Vacher et al. 2011, Xu, Zhong et al. 2013, Arnold and Gallo 2014, Ganguly, Tang et al. 2015). Cargo accumulation along such crowded paths appears to be a general feature of axonal transport arising from such common impediments to all types of transport.

Our experiments show that multiple types of vesicular cargo in different neurons and across different organisms indeed stall for times up to several minutes (Figure 1A-F, 6A). We observe that nearly 90% of long-lived stationary pre-SVs and majority of other types of cargo we examined that are stationary are present at actin-rich regions (Figure 1G, H, 6A-C, 2A-C). Such stalling can arise from the physical constraints caused by the actin mesh itself. In the case of *C. elegans,* for instance, it is known that a filamentous architecture, likely composed of actin, is present in proximity to the plasma membrane and connects microtubules to it (Cueva, Mulholland et al. 2007). The average length of these filaments is around 14±3nm (Cueva, Mulholland et al. 2007). Though the pore size of the mesh is not known, multiple such filaments can potentially constrain cargo motion. However, cargo that stall in actin-rich regions could also be captured by myosins present in such regions that then bind to the cargo (Pathak, Sepp et al. 2010, Bittins, Eichler et al. 2010). Cargo stalling is also likely to depend on a combination of cargo size, composition of the motors on the cargo surface, stiffness of the cargo and any actin binding proteins on the cargo surface.

Since different types of cargo stall at actin-rich locations (Figure 1G, H, 2A-C), we suggest that physical crowding itself provides a mechanism for such stalling that can operate independently of specific myosin-cargo interactions. However, our experiments do not exclude the role of myosins in cargo capture. We also observe that the presence of stalled cargo at actin-rich regions increases the propensity of other types of cargo to stall at the same location (Figure 4A, 6F). Again, such stalling independent of cargo-type can arise as a result of additional physical crowding induced by the presence of pre-existing stalled cargo at a location (Figure 4B, C).

The lack of cargo specificity is an argument in favour of physical effects dominating over motor-based ones. These cargo themselves vary in size from 30nm to 3μm (Tsukita and Ishikawa 1980, Nakata, Terada et al. 1998, Cueva, Mulholland et al. 2007) and thus can locally clog the axonal transport path leading to local traffic jams. This is expected in TRNs which have an average diameter of ~4μm that is filled with ~45, 15 protofilament microtubules with an outer diameter of ~30nm thus occupying a large volume of the available space (Chalfie and Thomson 1982).

Additionally, such physical crowding can itself influence motion properties through motors. A previous study shows that stalled cargo slow the motion of other moving cargo in close proximity (Che, Chowdary et al. 2016). Further the accumulation of motors at the ends of microtubules *in vitro* itself influences motion of motors walking along the tracks (Leduc, Padberg-Gehle et al. 2012). Our data show that motors can accumulate along with stationary vesicles likely at actin-rich regions which may similarly influence motion of vesicles at close proximity (Figure S6E). This enriched pool of motors can allow different cargo to sample multiple motors and perhaps alter their trajectory of motion. Thus cargo stalling might arise both through direct physical crowding and also via indirect and more complex effects mediated through motors. The net consequence of such effects is a local traffic jam and accumulation of cargo.

Given that multiple sites along the neuronal process are prone to traffic jams, processes that resolve these traffic jams to maintain cargo flow must exist. We observe that stationary cargo do mobilize resulting in a three-fold increase in cargo flow through the same location (Figure 5B, C). Cargo emerge from already existing stationary cargo at regular intervals (Figure 7C-E). Thus, stationary vesicular cargo clusters can function as dynamic reservoirs that can both trap and release vesicles. The mobilization of such vesicles could depend on the state of motors as well as subtle changes in cytoskeletal architecture. Our data suggests that levels of motor on the cargo surface affect both stalling and emergence of cargo from such locations (Figure S6F, G). Additionally, calcium influx is also known to influence both motor behavior and cargo motion in a neuron (Wang and Schwarz 2009, Hoerndli, Wang et al. 2015). Vesicle capture and release may be selectively modulated from stationary vesicle pools in response to Ca^2+^ concentration changes arising from external or internal stimuli. The ability of stationary pools of vesicles to function as dynamic vesicle reservoirs may thus provide an additional functional layer of control of cargo flux in response to cellular requirements, with large variations in flux potentially buffered by clusters releasing or trapping vesicles.

## MATERIALS AND METHODS

### Strains

Following strains were used in the study:

|  |  |  |
| --- | --- | --- |
| *jsIs37* | *p<sub>mec-7</sub>::SNB-1::GFP* (Nonet, 1999) | NM664 |
| *jsIs821* | *p<sub>mec-7</sub>::GFP::RAB-3* (Kumar, Choudhary et al. 2010) | NM2689 |
| *jsIs682* | *p<sub>rab-3</sub>::GFP::RAB-3* (Mahoney, Liu et al. 2006) | NM2415 |
| *wyIs291* | *p<sub>unc-86</sub>::GFP::utCH*; *p<sub>ODR-1</sub>::GFP* (Chia, Patel et al. 2012, Chia, Chen et al. 2014) |  |
| *twnEx195* | *p<sub>mec-7</sub>::COR-1::mCherry*, *p<sub>ttx-3</sub>::mCherry* | CLP945 |
| *tbIs263* | *p<sub>mec-7</sub>::COR-1::mCherry*, *p<sub>ttx-3</sub>::mCherry* | TT753 |
| *jsIs1073* | *p<sub>mec-7</sub>::tagRFP-MITO unc-119* (Zheng, Ahlawat et al. 2014) | NM2057 |
| *jsIs1239* | *[unc-119(+), p<sub>mec-7</sub>::DENDRA-2::RAB-3 RIM-3, unc-119]* | NM2357 |
| *oxSi266* | *unc-119(ed-9)*; *oxSi266[p<sub>rund-1</sub>::eGFP::RAB-5, unc-119]* (Ailion, Hannemann et al. 2014) | EG6193 |
| *tbIs227* | *[p<sub>mec-4</sub>::mcherry::RAB-3; p<sub>unc122</sub>::GFP]* | TT1884 |
| *tbIs147* | Pan neuronal UNC-104::GFP integrated in *unc-104(e1265)* background | TT343 |
| *rab-3(js49)* | complete loss of function allele that has a nonsense mutation in the tryptophan 76 codon, (Nonet, Staunton et al. 1997) | NM791 |
| *unc-104(e1265tb120)* | Suppressor of *unc-104(e1265)*, *UNC-104(D1497N M1540I)*, (Jitendra et al. 2010) | TT385 |

*jsIs1239* was created by inserting the pmec-7:: Dendra2-RAB-3 Rim3’sequences from plasmid NM2357 into chromosome II at the mos ttTi5605 insertion site using MosSCI in an *unc-119(ed3)* mutant background (Frokjaer-Jensen, Davis et al. 2008). The structure of insertion was confirmed by PCR and restriction digestion of the resulting PCR product.

NM2357 was constructed by PCR amplification of *dendra2* from pDendra2-C (Evrogen, Moscow Russia) using oligonucleotides

TTTAgctagcgtcgacggtacCATGAACACCCCGGGAATT and TTCATGTACACGCCGCTGTC, digesting the resulting product with NheI and BsrGI, and inserting into similarly digested NM2211 (pCFJ150-pmec-7::GFP::RAB-3 Rim3’) replacing GFP sequences with RAB-3.

NM2211 was created by inserting the pmec-7-GFP-RAB-3 seqeunces from as a BssHII/XbaI fragment from NM1028 (Bounoutas A. Zheng Q et al 2009) and inserting the fragment into BssHII/AvrII digested CFJ150.

The strain *mec-10(tm1552)* was confirmed using PCR with the following primer set: FP TGGGAGGGAGCTTCATCTTA RP GTAGGGTCTGCAACTAGCTC

### Worm maintenance

Worms were maintained on NGM agar plates seeded with OP50 *E. coli* strain, at a temperature of 20°C (Brenner 1974). L4s or 1 day adults from contamination-free and non-crowded plates were used for imaging.

#### *Drosophila* stocks

*Drosophila* stocks expressing Cha19bGal-4 UAS-GFP were raised on a standard corn meal medium at 25°C. Cha19bGal-4 drives the expression of UAS-RAB-4-mRFP and UAS-Lifeact-GFP specifically in cholinergic neurons of *Drosophila* (Salvaterra and Kitamoto 2001, Riedl, Crevenna et al. 2008). For dual color experiments and LatA treatment, filet prep was done on third instar larvae as described in Parton *et al.* 2010 and Brent *et al.* 2009.

### Dynamic imaging and analysis

#### Time-Lapse Imaging

Live single worms mounted on glass slides with agar pads were anesthetized using 5mM levamisole (Sigma-Aldrich, St. Louis, MO, USA) prepared in M9 buffer. For Latrunculin A injection, different concentrations of Latrunculin A (Sigma-Aldrich, St. Louis, MO, USA) solutions were prepared in 1x PBS. Latrunculin A or 1xPBS (as control) was injected into the pseudocoelom of L4 worms using a microinjection apparatus. The worms were anaesthetized using 5mM Levamisole immediately after injection and imaged.

Olympus IX83 microscope with spinning disc (Perkin Elmer Ultraview, Waltham, MA, USA) was used for imaging. Time-lapse fluorescence images of specific regions of the neuronal processes of TRNs were acquired at either 5fps (frames per second), or 3fps using a 60X objective of NA-1.63, and an Andor (iXon DU897-UVB, Belfast, UK)/Hamamatsu (SZK, Japan) monochrome camera. The typical length of neuron imaged is ~80μm in *C. elegans* TRNs and ~50μm in *Drosophila* chordotonal neurons.

#### Analysis

Kymographs were made using the plug-in MultipleKymograph built in ImageJ. Plugins were downloaded from the NIH website with the following links; http://www.rsbweb.nih.gov/ij/ and http://www.embl-heidelberg.de/eamnet/html/body_kymograph.html.

In the kymograph, cargo moving in the retrograde direction (towards the cell body), and anterograde direction (away from the cell body) were identified by sloped lines. Vertical lines represented stationary cargo. A cargo is counted as moving if it has been displaced by at least 3 pixels in successive time frames (Mondal, Ahlawat et al. 2011).

#### Motion properties calculations

Pause time is defined as the length of time for which a cargo stays stationary between two consecutive runs. Cargo pausing in proximity or at stationary cargo is not included in this analysis. The Measure option in ImageJ was used to measure pause time and the pixel value was multiplied by a conversion factor of either 0.2s (for 5fps) or 0.3s (for 3fps) depending on the frame rate of the acquired movie. Total run length is the sum of segment run lengths of a given vesicle. Segment run length is the distance moved by the cargo between pauses. Pause frequency refers to number of pauses per μm and is calculated by dividing number of pauses with the total run-length. For calculating motion properties we take into account all moving particles in a kymograph.

#### Cut offs for stationary pre-SVs

To calculate cut offs for stationary cargo, pause times were measured from kymographs. Maximum pause time for each cargo was quantified. Median value of maximum pause times quantified for all the cargo was used for these cut-offs for each imaged cargo. A cargo was defined as a stationary short-lived cargo if it stopped for at least five times the maximum observed pause time of that particular cargo (Table S2). Any cargo that is stationary for duration three times longer than the cut-off of a short-lived cargo was considered a long-lived stationary cargo.

#### Density calculation

To calculate the density of stationary cargo in a particular kymograph, several 20μm stretches in sharp focus along the neuronal process were selected. The numbers of long-lived stationary cargo were counted in each stretch. The density was represented as number of long-lived stationary cargo per 10μm of the axonal length.

#### Approximation of number of vesicles at a stationary cargo

We measure the intensities of vesicles when they pause between two consecutive run-lengths. For measuring intensities of both paused and stationary vesicles, we draw a line of thickness 2pixels spanning the stalled cargo. Measure option in ImageJ is used for quantifying the average intensities. Multiple such vesicles are sampled in a kymograph. We divide the intensity of a stationary cargo with the average intensity of paused vesicles to approximate the number of vesicles at a stationary cargo in the same kymograph.

#### Photobleaching and Photoconversion

Photoconversion experiments were performed with a spinning disc confocal system with an attached photokinesis unit (Perkin Elmer Ultraview, Waltham, USA). Time lapse images were acquired by using a 100X, 1.63 NA oil immersion, objective lens. Anesthetized animals were imaged along the neuronal process of PLM neurons. Spot function in software was used to randomly photoconvert stationary pre-SVs while a 5min movie was being acquired. Photoconversion of Dendra-2 was done using a 405nm LASER at 60% transmission for 100ms, 10 iterations of 10ms each. Kymographs were then used to analyze the events occurring at individual stationary pre-SVs.

#### Dual color imaging

Both simultaneous and sequential imaging of red and green markers was done for a total of 2 minutes in a Zeiss LSM 5 live scanning confocal system or Perkin Elmer spinning disc microscope using a 60X, oil immersion objective lens. Multispec beads were used to check the alignment of both cameras for dual color imaging. To check bleed through, laser power in one channel (e.g. green) was slowly increased and an increase in intensity in the other channel (e.g. red) was examined. This confirmation was done for all the fluorophore combinations used. Sequential or simultaneous imaging conditions were then accordingly chosen to collect unbiased data for a given pair of fluorophores depending on the bleed through. Time lapse images were acquired at a maximum frame rate of 1.5fps or 3fps. Alternate frames were separated using Image-J>> Stacks>> Tools>>Substack command. Kymographs were plotted for separated frames and overlaid for analysis.

#### Statistical analysis

Graph pad Prism 5.0 was used to perform the Student’s t-test or one-way ANOVA to compare between two groups or multiple groups respectively. Kruskal Wallis and Dunn’s comparison post tests were applied for Student t-test and one-way ANOVA respectively since not all the groups compared passed the normality tests. Two independent normality tests were used; D’Agostino and Perason omnibus normality test and Kolmogrov-Smirnov test (with DallalWilkinson-Lillie for P value). In case of comparison of flux with and without a stationary cargo at the same site, paired t-test was used. To test the statistical significance of data like stop and go or juxtaposition, represented as grouped plots in Figure 4 and Figure S6, percentage values from individual animals were compiled and tested for significance using the same statistical tests listed above. However for ease of representation, the errors bars are not included in the graphs and the data is represented as a sum of values from all individual animals. We have mentioned the p values in these cases in the legend.

Pearson correlation analysis was used to determine correlation between the density of stationary pre-SVs and run-length of moving cargo in a region. Correlation values were plotted on the graph and were fit using linear regression. p-values of < 0.001 is designated by ***, < 0.01 is designated by **, and < 0.05 is designated by *. All data are represented as mean±SEM in the graphs unless stated otherwise.

#### Statistical analysis of marker co-localization

We analyze snapshots obtained from multiple kymographs, about 7 per kymograph, from kymograph trajectories separated by at least 15s in time to ensure that they are approximately independent and across multiple worms (n=7-10). These yield spatial signals for the two fluorescent markers whose co-localization we wish to assess. We compute the Pearson’s Correlation Coefficient, the Mander’s Overlap Coefficient, and the Mander’s Correlation Coefficient, all measures of the correlations between the two markers. We use the Manders overlap coefficient (MOC) and the Mander’s correlation coefficients (MCC) to estimate marker colocalization. The MOC and the MCC should be most appropriate to our experimental situation, where we expect a strong overlap between signals from two markers only at isolated points along the axon, identified with actin accumulations, but very little overlap or correlation elsewhere, since at least one of these markers is associated with mobile cargo. We note that the MCC’s directly measure colocalization, since they compute the fraction of total probe fluorescence that colocalizes with the fluorescence of a second probe, irrespective of any linear relationship between their measured intensities. We also develop an approach based on the histogram of values of the scaled correlation function as a function of pixel location. For marker locations which are uncorrelated, the histogram should exhibit values equally distributed about zero, with vanishing skewness. If, on the other hand, these markers are correlated positively, we expect that the histogram will skew towards positive values. Thus an increased co-localization of the two markers is associated with a positive value of the mean and skewness of the histogram of correlation function values, which we examine using one-tailed t-tests of significance, computing the t-statistic and the Zg1 statistic from the data.

## ACKNOWLEDGEMENTS

We gratefully acknowledge support from DAE Project 12-R&D-IMS-5.02-0202 to GIM and SPK and HHMI. We thank Dr. Krishnamurthy CIFF-NCBS (No. SR/55/NM-36-2005) for access to imaging equipment, Sunaina Surana for LatA injections, Prof. Krishanu Ray for *Drosophila* stocks and lab space, Aparna Ashok for acquiring movies of commissures motor neurons in *C. elegans,* Madhushree Kamak for the schematic of *Drosophila* neuron. We thank all Koushika lab members for comments on the manuscript. Salary support KM: DBT post-doctoral fellowship, VK: IMSc Prism DAE. Research costs: To SPK: DST, CSIR, HHMI-IECS. GIM: DAE-SRC Fellowship and sabbatical support from the NUS, Singapore.

## REFERENCES

Ailion, M., M. Hannemann, S. Dalton, A. Pappas, S. Watanabe, J. Hegermann, Q. Liu, H. F. Han, M. Gu, M. Q. Goulding, N. Sasidharan, K. Schuske, P. Hullett, S. Eimer and E. M. Jorgensen (2014). “Two Rab2 interactors regulate dense-core vesicle maturation.” Neuron 82(1): 167–180.

Arnadottir, J., et al. (2011). “The DEG/ENaC protein MEC-10 regulates the transduction channel complex in Caenorhabditis elegans touch receptor neurons.” J Neurosci 31(35): 12695–12704.

Arnold, D. B. and G. Gallo (2014). “Structure meets function: actin filaments and myosin motors in the axon.” J Neurochem 129(2): 213–220.

Bálint, Š., I. Verdeny Vilanova, Á. Sandoval Álvarez and M. Lakadamyali (2013). “Correlative live-cell and superresolution microscopy reveals cargo transport dynamics at microtubule intersections.” Proc Natl Acad Sci U S A 110(9): 3375–3380.

Bittins, C. M., T. W. Eichler, J. A. Hammer, 3rd and H. H. Gerdes (2010). “Dominant-negative myosin Va impairs retrograde but not anterograde axonal transport of large dense core vesicles.” Cell Mol Neurobiol 30(3): 369–379.

Black, M. M. and J. T. Kurdyla (1983). “Microtubule-associated proteins of neurons.” J Cell Biol 97(4): 1020–1028.

Bounoutas A. Zheng Q. Michael L Nonet, Martin Chalfie (2009). “mec-15 Encodes an F-Box Protein Required for Touch Receptor Neuron Mechanosensation, Synapse Formation and Development”, Genetics 183(2): 607–617

Brenner, S. (1974). “The genetics of Caenorhabditis elegans.” Genetics 77(1): 71–94.

Burkel, B. M., G. von Dassow and W. M. Bement (2007). “Versatile fluorescent probes for actin filaments based on the actin-binding domain of utrophin.” Cell Motil Cytoskeleton 64(11): 822–832.

Cai, L., A. M. Makhov and J. E. Bear (2007). “F-actin binding is essential for coronin 1B function in vivo.” J Cell Sci 120(Pt 10): 1779–1790.

Chalfie, M. and J. N. Thomson (1979). “Organization of neuronal microtubules in the nematode Caenorhabditis elegans.” J Cell Biol 82(1): 278–289.

Chalfie, M. and J. N. Thomson (1982). “Structural and functional diversity in the neuronal microtubules of Caenorhabditis elegans.” J Cell Biol 93(1): 15–23.

Che, D. L., P. D. Chowdary and B. Cui (2016). “A close look at axonal transport: Cargos slow down when crossing stationary organelles.” Neurosci Lett 610: 110–116.

Chen, C. H., He C. W., Liao C. P., Pan, C. L. (2017). “A Wnt-planar polarity pathway instructs neurite branching by restricting F-actin assembly through endosomal signaling.” PLoS Genet 13(4): e1006720.

Chia, P. H., B. Chen, P. Li, M. K. Rosen and K. Shen (2014). “Local F-actin network links synapse formation and axon branching.” Cell 156(1-2): 208–220.

Chia, P. H., M. R. Patel and K. Shen (2012). “NAB-1 instructs synapse assembly by linking adhesion molecules and F-actin to active zone proteins.” Nat Neurosci 15(2): 234–242.

Coles, C. H. and F. Bradke (2015). “Coordinating neuronal actin-microtubule dynamics.” Curr Biol 25(15): R677–691.

Conway, L., D. Wood, E. Tüzel and J. L. Ross (2012). “Motor transport of self-assembled cargos in crowded environments.” Proc Natl Acad Sci U S A 109(51): 20814–20819.

Cueva, J. G., A. Mulholland and M. B. Goodman (2007). “Nanoscale organization of the MEC-4 DEG/ENaC sensory mechanotransduction channel in Caenorhabditis elegans touch receptor neurons.” J <u>Neurosci</u> 27(51): 14089–14098.

Dixit, R., J. L. Ross, Y. E. Goldman and E. L. Holzbaur (2008). “Differential regulation of dynein and kinesin motor proteins by tau.” Science 319(5866): 1086–1089.

Frokjaer-Jensen, C., M. W. Davis, C. E. Hopkins, B. J. Newman, J. M. Thummel, S. P. Olesen, M. Grunnet and E. M. Jorgensen (2008). “Single-copy insertion of transgenes in Caenorhabditis elegans.” Nat Genet 40(11): 1375–1383.

Ganguly, A., Y. Tang, L. Wang, K. Ladt, J. Loi, B. Dargent, C. Leterrier and S. Roy (2015). “A dynamic formin-dependent deep F-actin network in axons.” J Cell Biol 210(3): 401–417.

Gibbs, K. L., L. Greensmith and G. Schiavo (2015). “Regulation of Axonal Transport by Protein Kinases.” Trends Biochem Sci 40(10): 597–610.

Gu, Y., W. Sun, G. Wang, K. Jeftinija, S. Jeftinija and N. Fang (2012). “Rotational dynamics of cargos at pauses during axonal transport.” Nat Commun 3: 1030.

Hall, D. H. and E. M. Hedgecock (1991). “Ki nesi n-related gene unc-104 is required for axonal transport of synaptic vesicles in C. elegans.” Cell 65(5): 837–847.

Hirokawa, N. (1982). “Cross-linker system between neurofilaments, microtubules, and membranous organelles in frog axons revealed by the quick-freeze, deep-etching method.” J Cell Biol 94(1): 129–142.

Hirokawa, N., S. Niwa and Y. Tanaka (2010). “Molecular motors in neurons: transport mechanisms and roles in brain function, development, and disease.” Neuron 68(4): 610–638.

Hirokawa, N. and Y. Noda (2008). “Intracellular transport and kinesin superfamily proteins, KIFs: structure, function, and dynamics.” Physiol Rev 88(3): 1089–1118.

Hirokawa, N. and R. Takemura (2004). “Molecular motors in neuronal development, intracellular transport and diseases.” Curr Opin Neurobiol 14(5): 564–573.

Hirokawa, N. and R. Takemura (2005). “Molecular motors and mechanisms of directional transport in neurons.” Nat Rev Neurosci 6(3): 201–214.

Hoerndli, F. J., R. Wang, J. E. Mellem, A. Kallarackal, P. J. Brockie, C. Thacker, D. M. Madsen and A. V. Maricq (2015). “Neuronal Activity and CaMKII Regulate Kinesin-Mediated Transport of Synaptic AMPARs.” Neuron 86(2): 457–474.

Hong, K., et al. (2000). “In vivo structure-function analyses of Caenorhabditis elegans MEC-4, a candidate mechanosensory ion channel subunit.” J Neurosci 20(7): 2575–2588.

Iacobucci, G. J., N. A. Rahman, A. A. Valtueña, T. K. Nayak and S. Gunawardena (2014). “Spatial and temporal characteristics of normal and perturbed vesicle transport.” PLoS One 9(5): e97237.

Jaworski, J., C. C. Hoogenraad and A. Akhmanova (2008). “Microtubule plus-end tracking proteins in differentiated mammalian cells.” Int J Biochem Cell Biol 40(4): 619–637.

Kumar, J., Choudhary, B. C., Metpally, R., Zheng, Q., Nonet, M. L., Ramanathan, S., Klopfenstein, D. R. Koushika, S. P (2010). “The Caenorhabditis elegans Kinesin-3 motor UNC-104/KIF1A is degraded upon loss of specific binding to cargo.” PLoS Genet. 6(11): e1001200.

Kaasik, A., D. Safiulina, V. Choubey, M. Kuum, A. Zharkovsky and V. Veksler (2007). “Mitochondrial swelling impairs the transport of organelles in cerebellar granule neurons.” J Biol Chem 282(45): 3282–32826.

Kang, J. S., J. H. Tian, P. Y. Pan, P. Zald, C. Li, C. Deng and Z. H. Sheng (2008). “Docking of axonal mitochondria by syntaphilin controls their mobility and affects short-term facilitation.” Cell 132(1): 137–148.

Khare, S. M., Awasthi, A., Venkataraman, V., Koushika, S. P. (2015). “Colored polydimethylsiloxane micropillar arrays for high throughput measurements of forces applied by genetic model organisms.” Biomicrofluidics: 9(1):014111

Kumar, J., B. C. Choudhary, R. Metpally, Q. Zheng, M. L. Nonet, S. Ramanathan, D. R. Klopfenstein and S. P. Koushika (2010). “The Caenorhabditis elegans Kinesin-3 motor UNC-104/KIF1A is degraded upon loss of specific binding to cargo.” PLoS Genet 6(11): e1001200.

L, L. C. a. R. J. (2014). “Kinesin Motor Transport is Altered by Macromolecular Crowding and Transiently Associated Microtubule-Associated Proteins.” arXiv: 1409.345 5 [q-bio.BM].

Leduc, C., K. Padberg-Gehle, V. Varga, D. Helbing, S. Diez and J. Howard (2012). “Molecular crowding creates traffic jams of kinesin motors on microtubules.” Proc Natl Acad Sci U S A 109(16): 6100–6105.

Leterrier, C., H. Vacher, M. P. Fache, S. A. d’Ortoli, F. Castets, A. Autillo-Touati and B. Dargent (2011). “End-binding proteins EB3 and EB1 link microtubules to ankyrin G in the axon initial segment.” Proc Natl Acad Sci U S A 108(21): 8826–8831.

Maday, S., A. E. Twelvetrees, A. J. Moughamian and E. L. Holzbaur (2014). “Axonal Transport: Cargo-Specific Mechanisms of Motility and Regulation.” Neuron 84(2): 292–309.

Mahoney, T. R., Q. Liu, T. Itoh, S. Luo, G. Hadwiger, R. Vincent, Z. W. Wang, M. Fukuda and M. L. Nonet (2006). “Regulation of synaptic transmission by RAB-3 and RAB-27 in Caenorhabditis elegans.” Mol Biol Cell 17(6): 2617–2625.

Mondal, S., S. Ahlawat, K. Rau, V. Venkataraman and S. P. Koushika (2011). “Imaging in vivo neuronal transport in genetic model organisms using microfluidic devices.” Traffic 12(4): 372–385.

Nakata, T., S. Terada and N. Hirokawa (1998). “Visualization of the dynamics of synaptic vesicle and plasma membrane proteins in living axons.” J Cell Biol 140(3): 659–674.

Narayanareddy, B. R., S. Vartiainen, N. Hariri, D. K. O’Dowd and S. P. Gross (2014). “A biophysical analysis of mitochondrial movement: differences between transport in neuronal cell bodies versus processes.” Traffic 15(7): 762–771.

Nonet, M. L. (1999). “Visualization of synaptic specializations in live C. elegans with synaptic vesicle protein-GFP fusions.” J Neurosci Methods 89(1): 33–40

Nonet, M. L., J. E. Staunton, M. P. Kilgard, T. Fergestad, E. Hartwieg, H. R. Horvitz, E. M. Jorgensen and B. J. Meyer (1997). “Caenorhabditis elegans rab-3 mutant synapses exhibit impaired function and are partially depleted of vesicles.” J Neurosci 17(21): 8061–8073.

O’Hagan, R., et al. (2005). “The MEC-4 DEG/ENaC channel of Caenorhabditis elegans touch receptor neurons transduces mechanical signals.” Nat Neurosci 8(1): 43–50.

Pathak, D., K. J. Sepp and P. J. Hollenbeck (2010). “Evidence that myosin activity opposes microtubule-based axonal transport of mitochondria.” J Neurosci 30(26): 8984–8992.

Richardson, C. E., K. A. Spilker, J. G. Cueva, J. Perrino, M. B. Goodman and K. Shen (2014). “PTRN-1, a microtubule minus end-binding CAMSAP homolog, promotes microtubule function in Caenorhabditis elegans neurons.” Elife 3: e01498.

Riedl, J., A. H. Crevenna, K. Kessenbrock, J. H. Yu, D. Neukirchen, M. Bista, F. Bradke, D. Jenne, T. A. Holak, Z. Werb, M. Sixt and R. Wedlich-Soldner (2008). “Lifeact: a versatile marker to visualize F-actin.” Nat Methods 5(7): 605–607.

Ross, J. L., H. Shuman, E. L. Holzbaur and Y. E. Goldman (2008). “Kinesin and dynein-dynactin at intersecting microtubules: motor density affects dynein function.” Biophys J 94(8): 3115–3125.

Salvaterra, P. M. and T. Kitamoto (2001). “Drosophila cholinergic neurons and processes visualized with Gal4/UAS-GFP.” Brain Res Gene Expr Patterns 1(1): 73–82.

Shen, K., R. D. Fetter and C. I. Bargmann (2004). “Synaptic specificity is generated by the synaptic guidepost protein SYG-2 and its receptor, SYG-1.” Cell 116(6): 869–881.

Sheng, Z. H. (2014). “Mitochondrial trafficking and anchoring in neurons: New insight and implications.” J Cell Biol 204(7): 1087–1098.

Suzuki, H., et al. (2003). “In vivo imaging of C. elegans mechanosensory neurons demonstrates a specific role for the MEC-4 channel in the process of gentle touch sensation.” Neuron 39(6): 1005–1017.

Chatzigeorgiou, M., et al. (2010). “Spatial asymmetry in the mechanosensory phenotypes of the C. elegans DEG/ENaC gene mec-10.” J Neurophysiol 104(6): 3334–3344.

Tang, Y., D. Scott, U. Das, D. Gitler, A. Ganguly and S. Roy (2013). “Fast vesicle transport is required for the slow axonal transport of synapsin.” J Neurosci 33(39): 15362–15375.

Tang, Y., D. A. Scott, U. Das, S. D. Edland, K. Radomski, E. H. Koo and S. Roy (2012). “Early and selective impairments in axonal transport kinetics of synaptic cargoes induced by soluble amyloid β-protein oligomers.” Traffic 13(5): 681–693.

Tsukita, S. and H. Ishikawa (1980). “The movement of membranous organelles in axons. Electron microscopic identification of anterogradely and retrogradely transported organelles.” J Cell Biol 84(3): 513–530.

Vale, R. D. (2003). “The molecular motor toolbox for intracellular transport.” Cell 112(4): 467–480.

Vershinin, M., B. C. Carter, D. S. Razafsky, S. J. King and S. P. Gross (2007). “Multiple-motor based transport and its regulation by Tau.” Proc Natl Acad Sci U S A 104(1): 87–92.

Wang, X. and T. L. Schwarz (2009). “The mechanism of Ca2+ -dependent regulation of kinesin-mediated mitochondrial motility.” Cell 136(1): 163–174.

Wang, X., F. Zhou, S. Lv, P. Yi, Z. Zhu, Y. Yang, G. Feng, W. Li and G. Ou (2013). “Transmembrane protein MIG-13 links the W nt signaling and Hox genes to the cell polarity in neuronal migration.” Proc Natl Acad Sci U S A 110(27): 11175–11180.

Xu, K., G. Zhong and X. Zhuang (2013). “Actin, spectrin, and associated proteins form a periodic cytoskeletal structure in axons.” Science 339(6118): 452–456.

Yau, K. W., S. F. van Beuningen, I. Cunha-Ferreira, B. M. Cloin, E. Y. van Battum, L. Will, P. Schätzle, R. P. Tas, J. van Krugten, E. A. Katrukha, K. Jiang, P. S. Wulf, M. Mikhaylova, M. Harterink, R. J. Pasterkamp, A. Akhmanova, L. C. Kapitein and C. C. Hoogenraad (2014). “Microtubule minus-end binding protein CAMSAP2 controls axon specification and dendrite development.” Neuron 82(5): 1058–1073.

Yogev, S., R. Cooper, R. Fetter, M. Horowitz and K. Shen (2016). “Microtubule Organization Determines Axonal Transport Dynamics.” Neuron 92(2): 449–460.

Zheng, Q., S. Ahlawat, A. Schaefer, T. Mahoney, S. P. Koushika and M. L. Nonet (2014). “The vesicle protein SAM-4 regulates the processivity of synaptic vesicle transport.” PLoS Genet 10(10): e1004644.

Zhang, W., et al. (2008). “Intersubunit interactions between mutant DEG/ENaCs induce synthetic neurotoxicity.” Cell Death Differ 15(11): 1794–1803.

